# Relative Index of Chimeric Expression (RICE) Analysis: A Quantitative Approach for Chimeric RNAs Using *FusionBlaster*

**DOI:** 10.64898/2026.04.15.718257

**Authors:** Samuel Haddox, Yuhang Mao, Anam Tajammal, Jack Engel, Sarah Lynch, Ningxi Huang, Khalid Raby, Andrea Kian, Hui Li

## Abstract

Chimeric RNA molecules, which contain nucleotide sequences originating from multiple genes, are generated by chromosomal rearrangements, transcriptional read-throughs, or trans-splicing between separate parental transcripts. Chimeric RNAs have been functionally validated in both pathological and normal healthy physiological contexts indicating the biological significance of chimeric RNA expression. There is, however, currently no standard for computationally quantifying chimeric RNA expression and only limited benchmarking data available for the few chimeric RNA detection software that attempt to measure the abundance of the predicted chimeras. Here, we develop the relative index of chimeric expression, RICE, that is calculated based on the relative expression of chimeric transcripts compared to the respective parental WT transcripts. We evaluate three different methods for generating this measurement from simulated RNA sequencing data with known transcript abundances. Our BLAST-based approach outperforms STAR and Kallisto based approaches when considering both accuracy and consistency between simulated data of different read lengths and sequencing depths. We further demonstrate that RICE values can be validated using qPCR and are sensitive to dynamic conditions using siRNA targeting chimeric RNA expression. Finally, we apply our RICE analysis pipeline to clinical prostate cancer data. We quantify over 1200 chimeric RNAs in primary prostate cancer, metastatic prostate cancer, and non-cancer tissue samples from GTEx. Our differential RICE analysis revealed a clustering of prostate cancer tissue samples from three different sequencing cohorts <u>distinct</u> from their associated tissue type noncancer GTEx clusters. Our pipeline is publicly available on github and can be run on a personal laptop with computational resources and processing time dependent on the number of quantified chimeras.

## Introduction

Chimeric RNA molecules, which contain nucleotide sequences originating from multiple genes, are generated by chromosomal rearrangements, transcriptional read-throughs, or trans-splicing between separate parental transcripts (1). While it may be possible to generate chimeric artifacts during reverse transcriptase (RT) reactions in the lab, genuine chimeric RNAs have been functionally validated, correlated to phenotypes, and ultimately found as biologically relevant (2, 3). Since the discovery of BCR-ABL from the Philadelphia chromosome of chronic myelogenous leukemia patients in 1960 (4–6), gene fusions resulting in the transcription of chimeric RNAs have been associated with cancer. Indeed, we continue to find novel chimeric RNAs associated with various oncogenic phenotypes (7–11). However, it is now believed that chimeric RNAs are not only involved in the pathophysiology of cancer but also have non-pathological implications in normal physiological and developmental processes. This is supported by detection of several thousands of recurrent chimeric RNAs found to be actively transcribed through-out 41 different tissue types from over 500 noncancer individuals (1, 12, 13). The continued discovery of functional chimeric RNAs to date suggests that chimeric RNA synthesis expands the functional transcriptome (14).

The use of RNA sequencing (RNA-Seq) has played a critical role in the discovery of novel chimeric RNAs by enabling the high-throughput screening of samples for sequencing reads that fragmentally align to nonadjacent regions of the genome. While new sequencing technologies have been developed to allow long read direct RNA sequencing with the potential to overcome obstacles associated with the short read sequencing of cDNA (15), the majority of existing RNA-seq data was produced using library preparation kits with a RT reaction step and sequenced with short read technology, consisting of paired-end (PE) or single-end (SE) reads with sequencing lengths typically ranging from 37-150 base pairs (bps) and up to 300 bps. It is for this reason that short read RNA-seq remains a valuable resource for chimeric RNA discovery and functional association with observed phenotypes.

Chimeric-RNA prediction software for short-read RNA-seq, such as EricScript, STAR-Fusion, Pizzly, FuSeq, and SOAPfuse, rely on identifying single read alignments that cover a novel intergenic splice junction (junction reads) and discordant paired-end read alignments (spanning reads) that do not directly cover the junction but, nonetheless, suggest that a chimeric RNA transcript is present, as depicted in Figure 1-A(16–19). These tools output similar information: a list of predicted chimeric RNAs within a sample, providing genomic positions for the last base pair of the first gene and the first base pair of the second gene, along with other features of the chimera. However, differences in alignment and prediction filtering methods result in varying final predictions between the software.

**Figure 1.**
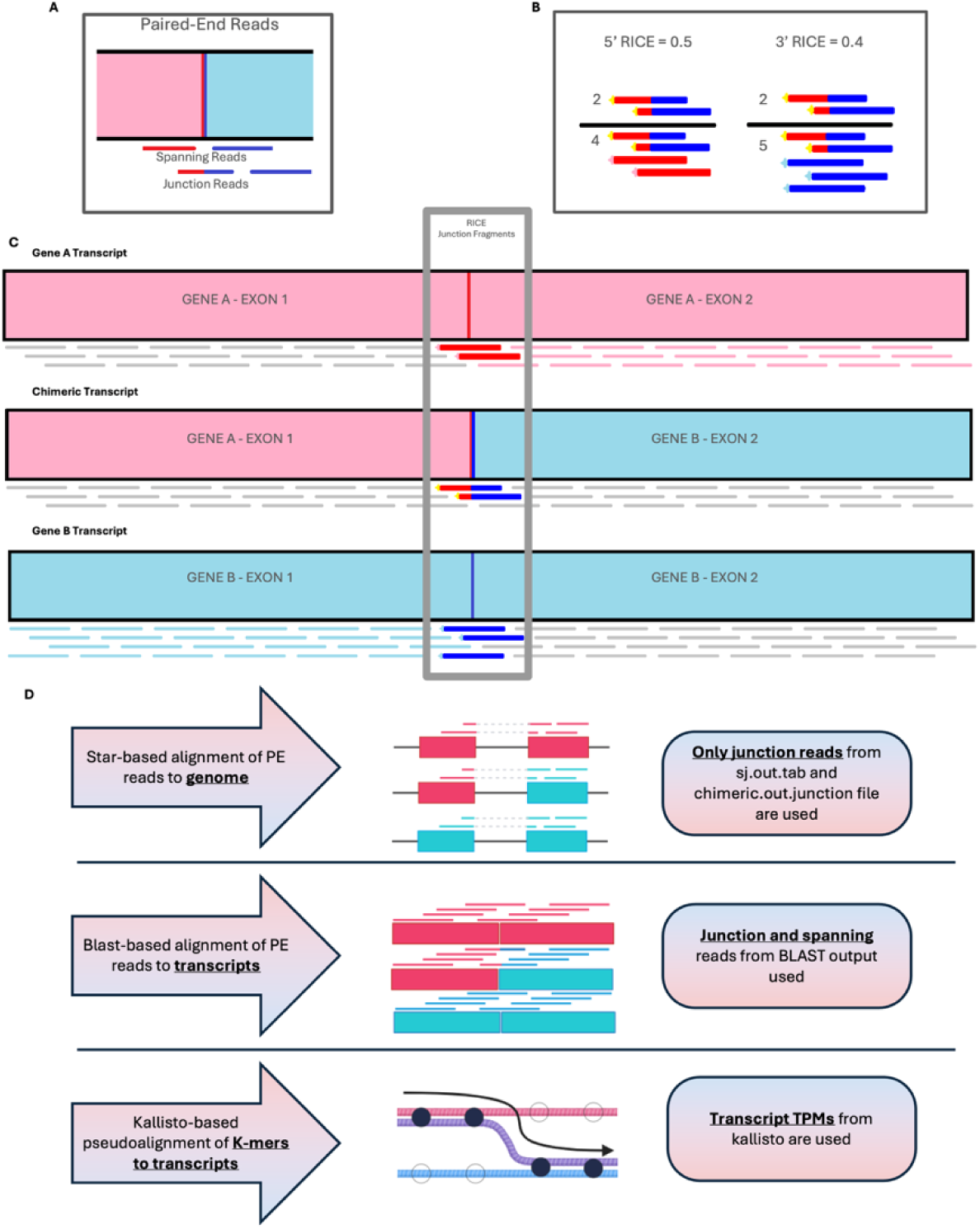
A) Junction reads are paired-end reads covering splice junction where one of the reads directly covers junction. Spanning reads are paired-end reads that straddling splice junctions. B) RICE is fraction of chimeric specific reads from total RICE junction reads. C) The RICE junction range is defined by library fragments that cover the chimeric specific junction and the respective WT parental splice junctions. D) Three approaches used to calculate RICE values includes: the STAR-based approach aligning reads to the genome, the BLAST-based approach where sequence reads are BLAST against a transcript reference, and a Kallisto-based approach directly using output transcript TPMs.

Many chimeric RNA detection software do not attempt to measure the expression level of the predicted chimeras. Of the chimeric RNA analysis tools, including Pizzly, EricScript, STAR-Fusion that attempt to measure the abundance of chimeric RNA transcripts or fragments, there is insufficient data on the accuracy of these quantification approaches. For example, STAR-Fusion includes a parameter for filtering chimeric predictions with fewer than 0.1 fragments per 10 million reads (FFPM), based on the 0.99 quantile of FFPMs for nonrecurrent chimeras in GTEx samples (16). However, this feature is not described as intended to provide an accurate quantification method and therefore no benchmarking data assessing the performance of star-fusion FFPM quantification is available. Pizzly and EricScript implement different quantification approaches, but their publications do not thoroughly evaluate the performance of these methods. Additionally, the Pizzly and EricScript approaches lack normalized quantification results, making it challenging to compare chimeric RNA expression across samples, tissue types, or studies with varying sequencing read lengths or depths.

In order to develop a method of relative chimeric RNA expression analysis, we considered a different conceptual approach that correlates more directly with the mechanisms that drive chimeric RNA generation rather than solely on their total transcript abundances. We also aimed to develop an approach that is directly comparable between different samples, tissue-types, sequencing conditions, and studies. For that reason, we have decided to calculate the chimeric transcript abundance as a fraction of the total abundance of transcripts containing the parent-chimera homologous region, hence known as the relative index of chimeric expression (RICE). This value differs from a traditional abundancy-based approach as it only considers reads that directly cover the chimeric junction and compares them to the amount of reads that cover the respective parental transcript specific junctions, as shown in Figure 1. The result of this calculation is the fraction of the chimeric transcripts from the total transcripts containing a region of the parental gene’s sequence. As such, each chimera has a 5’ RICE and a 3’ RICE that compares the chimera’s abundance to the 5’ parental gene and 3’ parental gene respectively.

We developed competing methods to make these calculations from PE RNA-Seq data based on alignment to the genome with STAR (Figure 1-D, top) and alignment directly to parental and chimeric transcripts with BLAST (Figure 1-D, middle). The STAR-based method is a simple but effective approach that only needs the junction read counts assigned to canonical splice sites from the sj.out.tab file and the chimeric junction read counts found in the chimeric.out.junction file. The BLAST-based method is more complex but captures both the junction and spanning reads, presumably allowing better sampling of chimeric and parental transcripts for calculation of RICE values. We then compared the STAR-based and BLAST-based methods to a quantitation approach that utilizes Transcript per Million (TPM) values generated by Kallisto (Figure 1-D, bottom), which is similar to the transcript specific quantitation approach employed by Pizzly for chimeric RNA quantitation (19).

In this study, we tested the accuracy of each approach on a simulated dataset with known chimeric RNA transcripts as well as known transcript abundances. To this end, we generated a simulated dataset based on the ChimPipe simulated data annotations (20). As the original dataset did not provide transcript abundances or any comment on relative expression between simulated samples, it was necessary to generate new simulated FASTQ files with known transcript abundances provided as TPM values. This data set consists of simulated samples of varying read length (75, 100, and 150) and varying library sizes (30, 40, 60, and 75 million reads) with relatively equal transcript abundances. We further validated the accuracy of our 2 approaches for calculating RICE values on HEK293T cell culture samples by comparing results from our computational analysis of RNA sequencing (RNA-seq) data with RICE values calculated using qPCR with standard curves. Finally, we demonstrated the utility of this approach by calculating RICE values for chimeric RNAs in clinical prostate cancer RNA-seq data.

## Methods

### Data Acquisition

RNA-seq data from normal human tissues were obtained from the Genotype-Tissue Expression (GTEx) project (V10 dbGaP Accession phs000424.v10.p2), comprising transcriptome profiles across 3 tissue types: prostate, liver, and lung (21–23). Primary prostate adenocarcinoma (PRAD) RNA-seq data generated by the Cancer Genome Atlas (TCGA) Research Network were downloaded via the Genomic Data Commons Data Portal (https://portal.gdc.cancer.gov/projects/TCGA-PRAD) (24). RNA-seq data for metastatic prostate cancer in Liver and Lung tissues samples were downloaded from National Institute of Health Sequence Read Archive (NIH SRA) from two projects GSE126078, SRA Accession SRP183532, and GSE118435, SRA Accession SRP157215 (22, 25).

### Data Simulation

Simulated paired-end FASTQ files with varying read lengths and read depths were generated based on the FASTA file containing the full length transcript sequences of chimeric and non-chimeric transcripts from the ChimPipe dataset (20); however, because the transcript abundancies were not specified for each transcript in the original ChimPipe dataset, simulated data was regenerated using the R package Rsubread function simReads (26).This allowed for a list of TPMs to be used as input to define known abundancies for each simulated transcript. The known TPMs for each transcript were used to calculate the simulated RICE values for benchmarking the STAR-Fusion-based, Kallisto-based, and BLAST-based (FusionBlaster) pipelines as shown below.

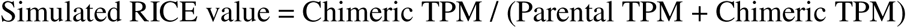

### Relative Index of Chimeric Expression Quantitation

RICE quantitation was performed using three different methods with simulated data to benchmark performance. The first method utilizes the SJ.out.tab and chimeric.out.junction files produced by the STAR aligner. The SJ.out.tab file is a tab separated values (tsv) formatted output that gives information about splice junction reads that cover canonical (non-chimeric) splice junctions. The chimeric.out.junction file is a tsv file that describes chimeric junction reads (16, 17). These files were merged allowing the junction read counts from both canonical and chimeric splice junctions to be counted and generate RICE values.

The second method of generating RICE values is based on a BLAST approach where the entire FASTQ files are converted to query FASTA files and blasted against a reference FASTA with the full length predicted chimeric transcript sequences as well as their respective parental WT transcripts. The full-length chimeric transcript sequences are predicted by taking all annotated 5’ parental WT transcript sequences upstream of the chimeric splice junction donor site and concatenating with all annotated 3’ parental WT transcript sequences downstream of the chimeric splice junction acceptor site to generate every possible pairwise combination. In order to optimize the performance of the BLAST we switched to using pBLAT, which allows parallel alignments to the reference index stored in memory (27). The BLAST formatted results are filtered for high confidence reads that specifically align to the chimeric or parental WT specific RICE junction window and stored in a data dictionary for rapid RICE value calculation.

We also tested an approach based on the pseudoaligner Kallisto. We utilized the full length predicted chimeric transcript sequences generated from the FusionBlaster reference building step in addition to the full ensembl annotation-based transcriptome as the reference for pseudoalignment. We then calculated RICE values using the Kallisto generated TPMs in the same fashion we calculated the simulated RICE values.

### Cell line culture

HCT116 and 293T cell lines were obtained from ATCC and cultured at 37°C/6% CO2 in Dulbecco’s Minimum Essential Medium (DMEM) supplemented with 10% fetal bovine serum (FBS), 1 mM sodium pyruvate, and 1% penicillin-streptomycin.

### qPCR Validation of RICE values of HEK293T cells

RNA was extracted from cultured HEK293T cells using the TRIzol reagent (Invitrogen). Total RNA was submitted to Novogene for PE-150bp RNA-Seq resulting in approximately 90 million reads. Chimeric RNAs were predicted using STAR-Fusion and cross referenced with chimeric RNAs previously validated in our lab and five chimeric RNAs were selected for qPCR validation. Primers were designed to span the chimeric junction, and to amplify the parental regions immediately upstream and downstream of the chimeric junction, as shown in Figure 4-A. All primers utilized for qPCR validation in HEK293T cells can be found in supplemental table 1. Only three of the five chimeric RNAs had specific amplification for all regions necessary to calculate RICE values. Standard curves were utilized to determine copy number and RICE values were determined as shown in Figure 4-C.

### siRNA and qPCR validation of SLC2A11-MIF RICE in HCT116

siRNAs (Invitrogen) were transfected using lipofectamine RNAiMAX (Invitrogen) according to the manufacturer’s instructions. Cells were plated in 6-well plates at 250,000 cells/well. At 24 hours post-plating and 70-80% cell confluency, cells were transfected with siRNAs. The non-targeting control siRNA sequence (siCT) used was CGUACGCGGAAUACUUCGA. The SLC2A11-MIF targeting siRNA (siSM) was UGCACCGCGAUGUAACUAA, which targets the junction of SLC2A11-MIF exon 8–exon 2. Cells were harvested for RNA isolation at 72 hours post-transfection. RNA was extracted using TRIzol Reagent (Invitrogen) followed by RNeasy mini (QIAGEN) kit. 1 µg of total RNA was sent to Novogene for PE-150bp sequencing and 1 µg of total RNA was transcribed into cDNA using Verso cDNA Synthesis Kit (Thermo) with random hexamers. A TaqMan assay was performed for SLC2A11-MIF quantitation to ensure chimeric specific amplification using 2x SensiFAST SYBR Hi-ROX (Bioline). RT-qPCR was performed for WT SLC2A11 and MIF with 2X Universal SYBR Green Fast qPCR Mix (ABclonal) using chimeric RNA exon-excluded gene specific primers (Eurofins Genomics). All primers and probes used for SLC2A11-MIF qPCR validation can be found in supplemental table 1.

### t-distributed Stochastic Neighbor Embedding

To investigate the distribution patterns of chimeric RNA signatures across different sample types, we performed dimensionality reduction and clustering analysis using t-distributed Stochastic Neighbor Embedding (t-SNE). The analysis incorporated RNA-seq data from multiple sources, including castration-resistant prostate cancer (CRPC) metastases to lung and liver (SRP183532, SRP157215), non-cancerous prostate, liver, and lung tissues from GTEx, and primary prostate adenocarcinoma (PRAD) samples from TCGA. After processing all RNA-seq datasets through our fusionBlaster pipeline with a reference database of 1,226 prostate cancer-associated chimeric RNAs, we extracted 5’ and 3’ RICE values for each chimeric RNA across all samples. These ratio values were combined to create a comprehensive feature matrix representing each sample’s chimeric RNA signature profile. We then applied t-SNE with two components and visualized the results with samples color-coded by tissue type and disease state.

### Motif and gene ontology analysis

Gapped Local Alignment of Motifs (GLAM2) tool from the MEME SUITE with its default settings was utilized to assess 5′ and 3′ parental genes located 200 bps upstream and downstream of the chimeric junctions to identify enriched motifs. Following this, the TomTom tool from MEME SUITE with default configurations and a database of known RNA binding protein motifs was used to identify the corresponding RNA binding proteins (RBP).

Additionally, the Gorilla tool was applied to predict gene ontology (GO) terms for the 5′ and 3′ parental genes of chimeras, using all annotated genes in GRChh38 as the background. Only the ten most significant hits were reported.

## Results

In order to produce a top performing method for differential RICE analysis we developed three different methods and benchmarked them using simulated data with known 5’ and 3’ RICE values. The R^2^ values from the Pearson Correlation analysis between each method and known values is shown in Figure 2-A, 5’ RICE values, and Figure 2-B, 3’ RICE values.

**Figure 2.**
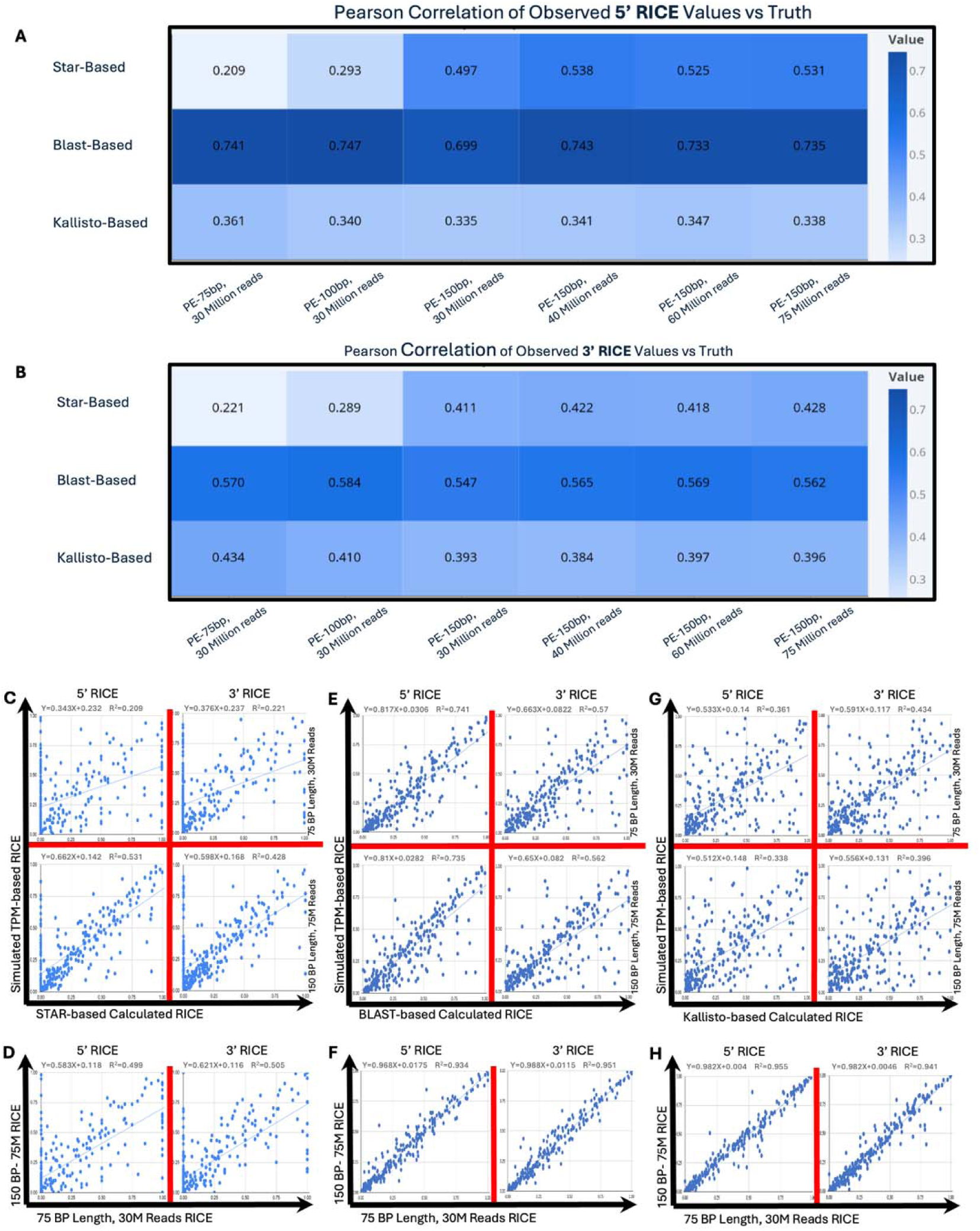
A) Pearson correlation R^2^ of known 5’ RICE values to calculated RICE values from three approaches. B) Pearson correlation of 3’ RICE values. C) STAR-based correlation plots showing accuracy of calculated RICE for 30 million reads at 75bp read length vs 75 million reads at 150bp read length D) STAR-based Correlation of plot showing consistency between 30 million reads at 75bp read length vs 75 million reads at 150bp read length. E) BLAST-based correlation plot for accuracy. F) BLAST-based correlation plot for consistency G) Kallisto-based correlation plot for accuracy. H) Kallisto-based correlation for consistency.

We first developed a method based on the STAR aligner that utilized the SJ.out.tab file output from the STAR aligner (17), which contains counts for all the splice junctions reads specific to the parental genes of chimeric RNAs, as well as the chimeric.out.junction file (16), which contains the splice junction reads for all chimeric RNAs. These files allowed for a simple and effective method to calculate 5’ and 3’ RICE values for all chimeric RNAs predicted by STAR in the simulated datasets, as shown in Figure 1-A,B,C and D using the number of chimeric junction specific reads divided by the sum of chimeric and parental specific reads. Due to the relatively small size of these two files, we were able to use a simple pipeline to calculate RICE values for STAR predicted chimeras using no additional computational resources than needed to run STAR. The time to run this pipeline only slightly increased the time of a standard STAR execution. This method performs relatively well at higher sequencing depth and longer sequencing read length, as shown in Figure 2-A, B, C and Supplemental Figure 1. However, due to the significantly worse performance at lower sequencing depth and shorter read length, results obtained by this method show a weaker correlation between simulated datasets as shown in Figure 2-D than the other methods utilized in this study. Additionally, STAR was not able to detect all of the simulated chimeras while using default parameters. Undetected Chimeric RNAs produced RICE values of 0, and consequently, the correlation between the calculated RICE values and known/true RICE was drastically reduced.

In order to determine the reasoning for the difference in performance of the STAR-based method between simulated datasets of varying read lengths, we investigated the differences in detectable chimeric read counts at different read lengths. We observed that the number of chimeric junction reads decreased and the number of chimeric spanning reads increased as read length was shortened. Junction reads and spanning reads are illustrated in Figure 1-C (16). Because the STAR-based approach solely relied on junction read counts, transcript sampling was reduced at shorter read lengths. We assessed the strength of the correlation between chimeric junction read counts between 75 basepair read length data and 150bp read length data and found a strong correlation with R^2^ =0.87, supplemental Figure 2-A. While the correlation between spanning read counts was much weaker, supplemental Figure 2-B, the correlation between total chimeric RNA reads, both junction and spanning, demonstrated the highest correlation with R^2^ =0.92, supplemental Figure 2-C. We reasoned that developing a method that was capable of utilizing both splice junction and spanning reads for sampling of the chimeras and their parental transcripts would produce more accurate RICE values.

To utilize both junction and spanning reads in the calculation of RICE values, we developed a pipeline that generates a reference fasta file consisting of chimeric and parental transcripts based on Ensembl transcript annotations (Figure 3-A) and performs a local alignment of RNA-Seq reads to the reference with pBLAT to detect RICE junction and spanning reads as depicted in Figure 3-B (27, 28). This pipeline is illustrated in Figure 3-C. While this pipeline only modestly outperforms the STAR-Fusion based approach at higher read depths and longer reads, the performance gains are much more notable at lower read depths and shorter reads (Figure 2-A, B, E, and supplemental Figure 3. The benefit of sampling chimeric and parental transcripts by both junction and spanning reads becomes more apparent when comparing the correlation between simulated data with lower reads depths and shorter reads to the dataset with higher read depth and longer reads. As shown in Figure 2-F, The FusionBlaster pipeline shows markedly increased consistency between RICE values obtained from the different simulated datasets.

**Figure 3.**
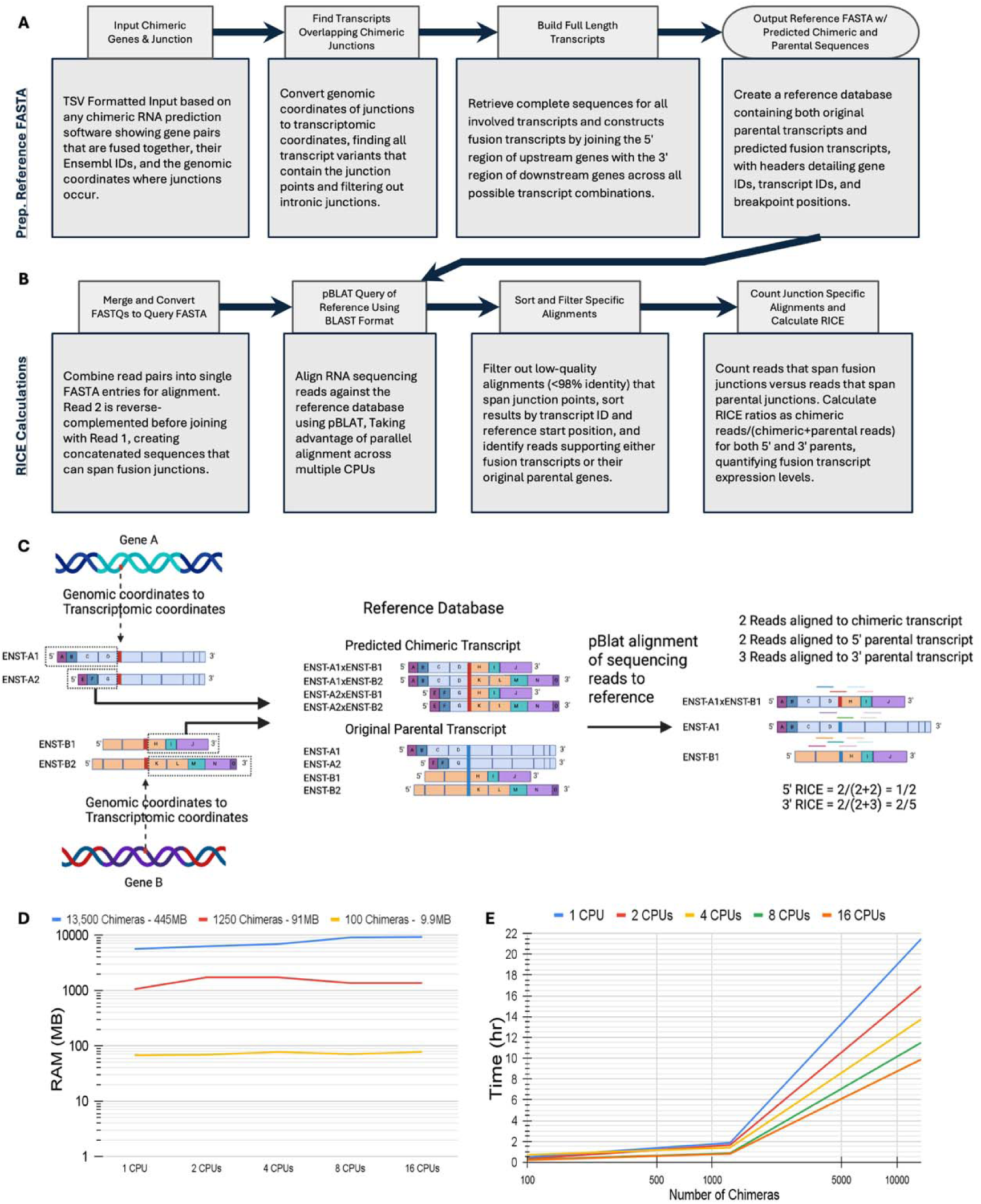
A) FusionBlaster pipeline details for reference FASTA preparation B) Detailed steps for FusionBlaster’s BLAT-based RICE calculation. C) Graphical representation of the FusionBlaster Pipeline. D) Memory requirements of FusionBlaster depending on the amount of Chimeras quantified. E) FusionBlaster processing time depending on number of cores used and chimeras quantified.

We then decided to benchmark a third approach based on a chimeric RNA quantification method in existing literature. Pizzly is an available fusion RNA detection and quantification software package that utilizes the pseudoaligner software, Kallisto (19) (29). Pizzly uses Kallisto to detect discordantly aligned reads as candidates for gene fusions. Transcript sequences for the gene fusions are then added to the transcriptome reference and Kallisto alignment is performed again to generate TPM values for the gene fusions, as well as all canonical transcripts. Based on this logic, we decided to utilize the same chimeric transcripts generated by the reference building step of the FusionBlaster pipeline with the Kallisto pseudoaligner to generate RICE values directly from the output TPMs in the same fashion as we calculated the known simulated RICE values. The Kallisto-based approach did not reproduce the simulated RICE values accurately, as shown in Figure 2-A,B,G and supplemental Figure 4; however, it is very interesting that this method did generate very consistent results across the simulated datasets with varying read depth and length (Figure 2-H).

The FusionBlaster pipeline outperformed the STAR-Fusion-based and Kallisto-based pipelines when considering both accuracy and consistency of reflecting simulated RICE values between datasets with sequencing depth ranging from 30 to 75 million reads and read lengths ranging from 75 to 150 base pairs. Because FusionBlaster is not a chimeric RNA detection software, it requires a list of chimeric RNAs as input for quantitation and can be easily implemented into a pipeline with any chimeric RNA detection software. FusionBlaster works with single or paired-end reads and has low computational resource requirements that scale with the number of chimeric RNAs being quantitated. FusionBlaster requires approximately 100MB of ram per 100 chimeric RNAs per sample (Figure 3-D) and can process 100 chimeric RNA quantitations in approximately 30 minutes or 1000 in less than 2 hours (Figure 3-E).

Next, we sought to validate FusionBlaster RICE values with quantitative real-time PCR (qPCR). We prepped RNA from HEK293T cells and sent for RNA-Seq using the Illumina platform, PE-150 basepair read length, and approximately 90 million reads. We predicted chimeric RNAs using STAR-Fusion and cross referenced those predictions with chimeric RNAs previously validated by PCR in our lab resulting in five chimeric RNA candidates (data not shown). We designed primers according to the scheme illustrated in Figure 4-A, where a primer pair spanning the chimeric junction specifically amplifies the chimeric transcript, and pairs upstream and downstream of the breakpoint point that do not span any isoform specific splice junction will amplify chimeric and parental transcripts together. For three out of the five chimeric RNA candidates, we were able to amplify the 5’, 3’, and chimeric templates specifically and at the correct amplicon length: POLA2-CDC42EP2(PC), DMC1-DDX19(DD), and BOLA-SMG1P6(BS). Using standard curves, Figure 4-B, we were able to determine the amount and copy number for the 5’,3’, and chimeric templates for PC, DD, and BS transcripts. Using the template amount and length we were able to calculate RICE values as shown in Figure 4-C. Figure 4-D, an agarose gel image, demonstrates that the standard curve qPCR amplicons were verified for specificity and expected length. When comparing the qPCR derived RICE values to the FusionBlaster calculated RICE values, we were able to obtain an average RICE accuracy of 92.3% (Figure 4-E). The FusionBlaster 3’ RICE values were more representative of the qPCR RICE values with an average accuracy of 96.1% compared to the 5’ RICE values that had an average accuracy of 88.5%.

We further validated FusionBlaster’s capacity for detecting changes in chimeric RNA expression using HCT116, a cell line known to express the chimeric RNA SLC2A11-MIF (SM) (10). Small interfering RNA (siRNA) molecules were designed for specific knock down of SM, SLC2A11 and MIF transcripts. Knockdown of SM (siSM) and parental transcripts (siSLC2A11 and siMIF) was performed alongside of a non-targeting control siRNA (siCTRL) in HCT116 before extracting RNA. Total extracted RNA was used to generate cDNA for qPCR and sent for RNA-Seq. FusionBlaster was used to determine the 5’ RICE (Figure 4-F) and 3’ RICE (Figure 4-G) values for each Knockdown. We performed qPCR on each sample by amplifying the chimeric RNA and quantifying the relative expression by using the 5’ parental gene as the endogenous control target (Figure 4-H), and the 3’ parental gene as the endogenous control target (Figure 4-I). FusionBlaster was able to detect the reduction in both the 5’ and 3’ Rice values from the siSM sample compared to the siCTRL sample. The reduction in both the 5’ and 3’ RICE values from the siSM sample was validated by qPCR. Figure # displays all significant P values as well as comparisons expected to yield significant changes in RICE values.

Finally, we employed FusionBlaster to quantitate over 1300 chimeric RNAs in clinical prostate cancer samples. Out of the 1350 chimeric RNAs detected in CCLE prostate cell lines, 1226 chimeric RNAs had sufficient parental transcript annotations for building a FusionBlaster reference FASTA and could be quantitated by the FusionBlaster pipeline. Using the reference FASTA containing the 1226 chimeric transcripts and their respective parental transcript sequences, we evaluated four separate RNA-seq datasets together. We took advantage of GTEx for non prostate cancer patient prostate, lung, and liver tissue samples. For prostate cancer patients we incorporated primary prostate cancer samples from TCGA with liver and lung metastatic prostate cancer samples from (22, 25) GSE126078 and GSE118435. The generalized workflow for our differential RICE analysis in prostate cancer is shown in Figure 5-A.

**Figure 4.**
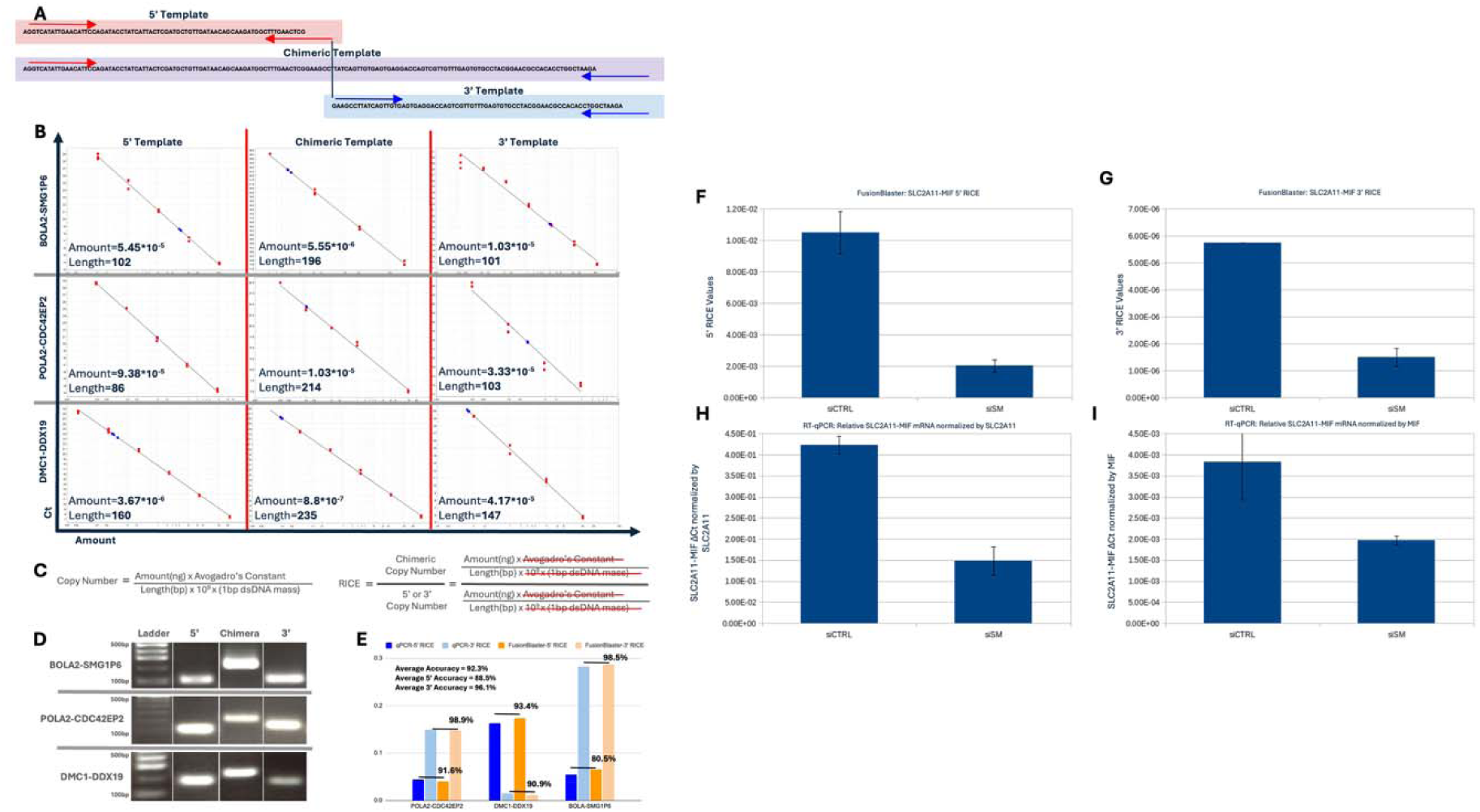
**A)** Primer design schematic for validating RICE values with RT-qPCR. **B)** Standard Curves used to calculate amount of target template in sample. **C)** Copy number Equation used for generating RICE value. **D)** Agarose Gel Electrophoresis confirming amplicon size. **E)** Accuracy assessment of FusionBlaster RICE values using RT-qPCR RICE values. **F, G)** FusionBlaster derived SLC2A11-MIF RICE values. **H, I)** RT-qPCR derived relative expression of SLC2A11-MIF as normalized by parental-WT genes.

**Figure 5.**
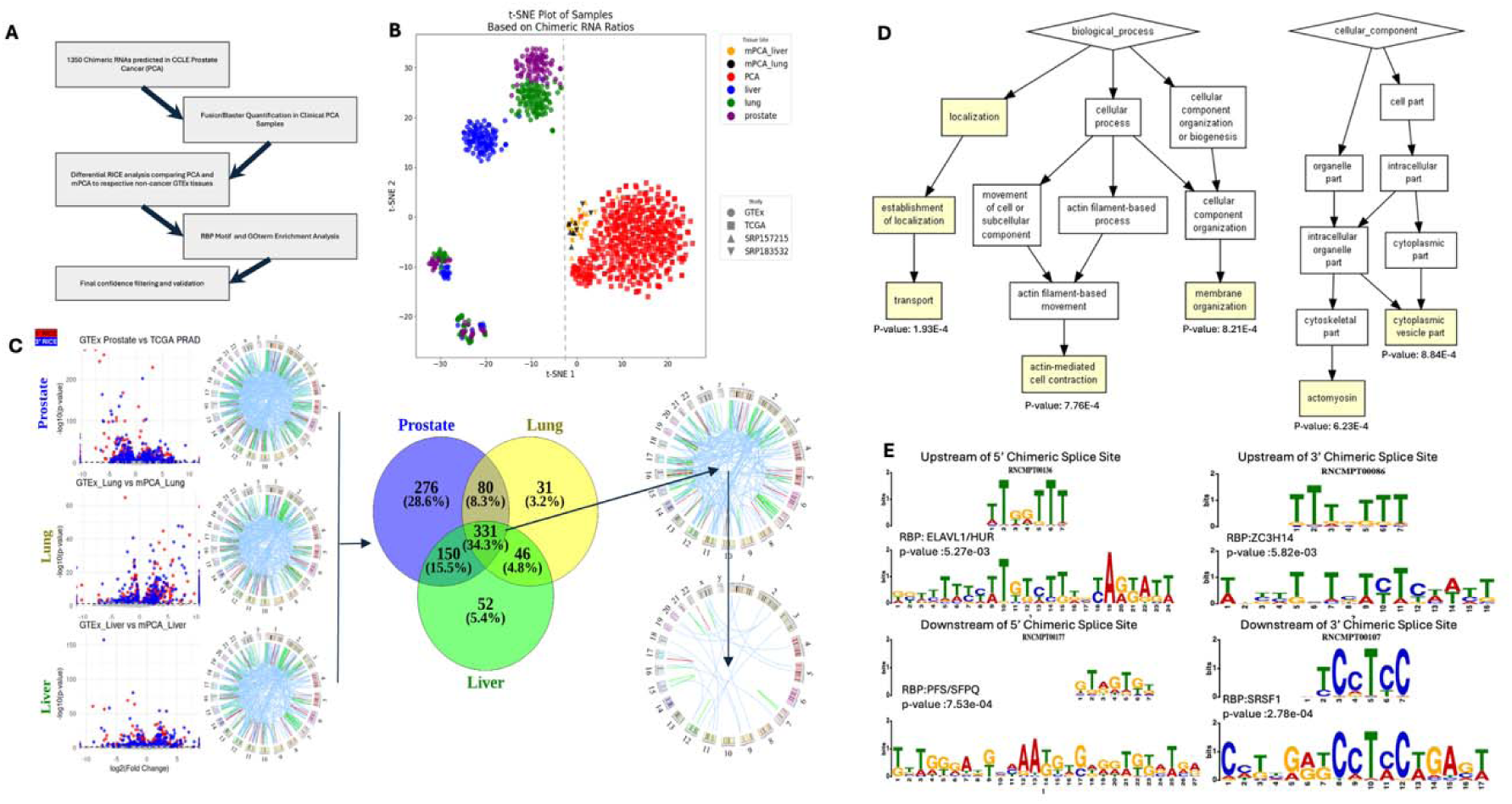
**A)** Flow chart of Prostate Cancer differential RICE analysis using FusionBlaster. **B)** tSNE plot of prostate cancer and non-diseased liver, lung, and prostate tissue clustering based on RICE profiles. **C)** Left, results of differential RICE analysis comparing each prostate cancer tissue type to its respective non-cancer tissue (5’ RICE - Red, 3’ RICE - Blue); Middle, colocalizing results produced 331 chimeric RNAs; Right, 28 out of 331 significantly different expressed chimeras have increased 5’ and 3’ RICE values. **D)** Biological processes and cellular components results from the GOrilla GO-term enrichment analysis of parental genes for 311 significantly different chimeras. **E)** RNA binding protein motif enrichment analysis using 200 bp upstream of 5’ chimeric splice site (Top Left), downstream of 5’ chimeric splice site (Bottom Left), upstream of 3’ chimeric splice site (Top Right), and downpstream of 3’ chimeric splice site (Bottom Right).

First, we performed tSNE clustering analysis on principal components derived from the fusionBlaster-based RICE values to demonstrate that differential RICE analysis is capable of distinguishing prostate cancer samples from noncancer patient controls, as seen in Figure 5-B. Our clustering analysis highlights the usefulness of FusionBlaster derived RICE values that are less influenced by technical sequencing factors as they are able to group primary prostate cancer from the TCGA cohort with metastatic prostate cancer in liver from (22, 25) GSE126078 and GSE118435 and metastatic prostate cancer in lung from (22, 25) GSE126078 and GSE118435.

Our tSNE clustering analysis illustrates a chimeric RNA expression signature associated with prostate cancer.

To deduce the chimeric RNA transcripts driving the prostate cancer chimeric RNA expression signature, we implemented Wilcoxon Ranked Sum into our differential RICE analysis to compare each prostate cancer tissue type, primary prostate, liver metastasis, and lung metastasis, against the respective noncancer tissue type from GTEx. As shown in Figure #-C, this resulted in 837 chimeric RNAs differentially expressed in primary prostate cancer vs GTEx prostate tissue samples (supplemental file), 579 chimeric RNAs differentially expressed chimeric RNAs in prostate cancer metastasis in liver vs GTEx liver samples (supplemental file), and 488 chimeric RNAs differentially expressed in prostate cancer metastasis in lung vs GTEx lung samples (supplemental file). Colocalizing the differentially expressed chimeric RNAs from the analysis of the three tissue types revealed a core set of 331 chimeric RNAs with significantly different 5’ and/or 3’ RICE values (supplemental file), which were further investigated for GOterm enrichment and RNA binding motif enrichment analysis. We narrowed down the differentially expressed chimeric RNAs further by filtering the core set of 331 chimeric RNAs down to those that contained both 5’ and 3’ RICE values significantly upregulated in prostate cancer samples from all three tissue types resulting in 28 chimeric RNAs that have potential to serve as prostate cancer biomarkers (Table 1).

**Table 1.** Chimeras with significantly increased 5’ and 3’ RICE values in Prostate Cancer.

| Chimera | 5' Gene Ensembl ID | 3' Gene Ensembl ID | 5' Chimeric Splice Site | 3' Chimeric Splice Site |
| --- | --- | --- | --- | --- |
| MT1A--MT2A | ENSG00000205362 | ENSG00000125148 | 16:56639936:+ | 16:56609263:+ |
| MMP24-AS1--NR6A1 | ENSG00000126005 | ENSG00000148200 | 20:35272005:- | 9:124519896:- |
| SLC1A5--FN1 | ENSG00000105281 | ENSG00000115414 | 19:46775053:- | 2:215431882:- |
| FAM76A--AACS | ENSG00000009780 | ENSG00000081760 | 1:27760073:+ | 12:125126581:+ |
| CD68--MPDU1 | ENSG00000129226 | ENSG00000129255 | 17:7582113:+ | 17:7583918:+ |
| AP5S1--LRIG2 | ENSG00000125843 | ENSG00000198799 | 20:3828837:+ | 1:113127270:+ |
| M6PR--MTRNR2L12 | ENSG00000003056 | ENSG00000269028 | 12:8941719:- | 3:96617621:- |
| CDK10--TBXAS1 | ENSG00000185324 | ENSG00000059377 | 16:89687690:+ | 7:139789372:+ |
| CD68--MPDU1 | ENSG00000129226 | ENSG00000129255 | 17:7582113:+ | 17:7583616:+ |
| NUPR2--SEPT7 | ENSG00000185290 | ENSG00000122545 | 7:56116015:- | 7:35872666:+ |
| CIAO1--SIK2 | ENSG00000144021 | ENSG00000170145 | 2:96273349:+ | 11:111728347:+ |
| TMPRSS2--ERG | ENSG00000184012 | ENSG00000157554 | 21:41498119:- | 21:38445621:- |
| CENPI--MREG | ENSG00000102384 | ENSG00000118242 | X:101163681:+ | 2:215944577:- |
| RCC1--C1QTNF3-AMACR | ENSG00000180198 | ENSG00000273294 | 1:28508160:+ | 5:34124528:- |
| SMAD2--FCN1 | ENSG00000175387 | ENSG00000085265 | 18:47827454:- | 9:134905966:- |
| LLGL2--AACS | ENSG00000073350 | ENSG00000081760 | 17:75564598:+ | 12:125126645:+ |
| PQBP1--SLC35A2 | ENSG00000102103 | ENSG00000102100 | X:48903143:+ | X:48903597:- |
| GTSE1--TRMU | ENSG00000075218 | ENSG00000100416 | 22:46330810:+ | 22:46330875:+ |
| SARNP--GDF11 | ENSG00000205323 | ENSG00000135414 | 12:55792948:- | 12:55753525:+ |
| CENPI--MREG | ENSG00000102384 | ENSG00000118242 | X:101163681:+ | 2:215944576:- |
| TMX2--RPS27L | ENSG00000213593 | ENSG00000185088 | 11:57740973:+ | 15:63151239:- |
| ZFYVE16--SLC9A7 | ENSG00000039319 | ENSG00000065923 | 5:80455431:+ | X:46604686:- |
| COX18--CBR3-AS1 | ENSG00000163626 | ENSG00000236830 | 4:73052835:- | 21:36175815:- |
| CHM--SPATA6L | ENSG00000188419 | ENSG00000106686 | X:85969739:- | 9:4627887:- |
| RABL2B--NUCB2 | ENSG00000079974 | ENSG00000070081 | 22:50770243:- | 11:17260590:+ |
| AKAP13--GOLGA8S | ENSG00000170776 | ENSG00000261739 | 15:85380798:+ | 15:23363678:+ |
| CD68--MPDU1 | ENSG00000129226 | ENSG00000129255 | 17:7582113:+ | 17:7583618:+ |
| RPL27--FOX L1 | ENSG00000131469 | ENSG00000176678 | 17:42998831:+ | 16:86578860:+ |

Using the GOrilla Gene Ontology suite, we performed GOterm enrichment analysis on the 331 prostate cancer core differentially expressed chimeric RNAs. As shown in Figure 5-D, we found that biological processes involved in transport, actin mediated cell contraction, and membrane organization were significantly enriched with respective p values of 1.93E-4, 7.76E-4, and 8.21E-4. Likewise, cellular components that were enriched were associated with the enriched biological processes and include actomyosin, with a p value of 6.23E-4, and cytoplasmic vesicles, with a p value of 8.84E-4. Actin mediated cell contraction and membrane organization processes via the actinomyosin cellular component are consistent with cancer metastasis and are not unexpected given the inclusion of metastatic prostate cancer samples in our analysis (32, 33)

Using upstream and downstream genomic sequences of 200 bp length from both the 5′ and 3′ chimeric junction sites as input for GLAM2, enriched sequences flanking the chimeric junctions of the core set of 331 differentially expressed prostate cancer chimeric RNAs were determined. The enriched sequences were then used as input for Tomtom to identify potential RNA binding protein (RBP) motifs associated with the enriched sequences. Several enriched sequences and RBP motifs were found (supplemental file); The highest scoring enriched RBP motifs and their associated enriched sequences are shown in Figure 5-E. ELAV-Like Protein 1 (ELAVL1) also known as Human Antigen R (HUR) (34), with a P-value of 5.27 × 10^−3^ was identified to have an RNA binding site motif found in sequences enriched in the upstream region of 5′ parental gene chimeric junction sites; while splicing factor proline and glutamine rich (SFPQ), with a *P*-value of 7.53 × 10^−4^ was identified to have an RNA binding site motif found in sequences enriched in the downstream region. Zinc Finger CCCH-Type Containing 14 (ZC3H14) (35), with a *P*-value of 5.82 × 10^−3^ was identified to have an RNA binding site motif found in sequences enriched in the upstream region of 3′ parental gene chimeric junction sites; Serine And Arginine Rich Splicing Factor 1 (SRSF1), with a *P*-value of 2.78 × 10^−4^ was identified to have an RNA binding site motif found in sequences enriched in the downstream region. All of the top enrichment RBP have previously been reported to be implicated in oncogenic pathologies; three out of the four RBPs, ELAV1, SFPQ, and SRSF1 have been specifically reported to play a role in prostate cancer progression and metastasis (36–38).

## Discussion

In this study we conceptualized a novel parameter for describing chimeric RNA expression, the relative index of chimeric expression (RICE). RICE values do not describe the actual abundances of chimeric RNA transcripts; instead, they define the chimeric portions of transcripts containing parental-chimera homologous sequences. RICE values can, however, be utilized to correct canonical transcript abundances by adjusting read counts based on the presence of a determined proportion of chimeric transcripts. RICE values also offer insight on the extent to which transcriptional read-through is happening at a given genomic locus and may also be useful in monitoring subpopulations of cells with genomic rearrangements that lead to the transcription of a gene fusion.

We developed three initial computational approaches, STAR-based, BLAST-based, and Kallisto-based pipelines capable of calculating RICE values from RNA-Seq data and benchmarked each using simulated datasets with varying read depth, read length, and known RICE values. While the STAR-based approach provides accurate RICE values with longer reads and higher read depth, this approach was limited to only calculating RICE values for chimeras that STAR captured reads for in the chimeric.out.junction file. Because the STAR-based approach demonstrated significantly increased performance at high read lengths and read depths, the resulting RICE values were not consistent between datasets of varying read depth and length. We believe the inconsistency between datasets is largely attributed to the STAR-based method relying solely on junction reads to calculate RICE values. Next, we applied a BLAST-based method capable of capturing both junction and spanning reads, allowing increased sampling of chimeric and parental reads. The BLAST-based method exhibited increased consistency across datasets of varying read depth and length. The BLAST-based method also demonstrated higher accuracy as measured by the Pearson correlation of calculated RICE values to true RICE. We then wanted to compare how well the pseudoaligner Kallisto would perform using the same chimeric transcripts references generated from the BLAST-based pipeline. While the resulting RICE values from the Kallisto-based approach were more accurate than the STAR-based approach at lower read lengths, BLAST and STAR based methods performed better on datasets with higher read lengths. It is interesting that even with less accurate RICE values the Kallisto-based method exhibited the highest degree of consistency between datasets with varying read depth and length. Overall, we considered the BLAST-based approach the best when considering both accuracy and consistency.

We then developed the BLAST-based approach into a streamlined, fast, and computationally resource efficient pipeline named FusionBlaster. The FusionBlaster pipeline replaces the original BLAST with pBLAT so that alignment of query sequences to the indexed reference sequences takes place in memory and is processed in parallel over a user specified number of processors. To keep memory requirements low, the pipeline splits query sequences into batches, each batch is then split across the specified number of processors during the BLAT step, combining the benefit of low memory serial batch-processing with the parallel processing of query sequences within a batch. The size of each batch is currently a pre-defined limit to allow the FusionBlaster pipeline to run on computers with limited memory; however, we expect to modify this parameter in future releases to be a user defined limit allowing users with access to computers with increased memory to run the pipeline with faster finishing times.

The FusionBlaster pipeline is dependent on transcript annotations for building full length chimeric RNA sequences from different combinations of the parental gene isoforms. While this was intended to allow alignment of paired end reads from larger RNA fragments, it creates additional use limitations and performance implications. Primarily, when the chimeric breakpoints do not fall within annotated exon boundaries, FusionBlaster cannot build a chimeric transcript reference sequence and thus the chimera cannot be quantified. Furthermore, the use of full-length chimeric transcripts in the BLAT reference fasta are not necessary as reads that do not fall within the RICE junction window are not considered for RICE value calculation. By implementing an approach to predict the distribution of RNA fragments in the sequencing library the length of reference transcripts can be optimized to only cover the necessary RICE junction window of parental and chimeric transcripts. While this approach would have the potential to speed up the BLAT step and downstream processing of the read alignments it does have the potential to miss RICE junction spanning reads with larger than expected insert sizes.

Validation of FusionBlaster RICE values can be achieved by several different approaches depending on the priority. In this study we demonstrate that the actual RICE value can be validated using qPCR with a standard curve to determine template copy number for RICE calculation. Using digital or digital droplet PCR may also be useful for validating RICE values more directly without the need of a standard curve. While digital PCR techniques may increase throughput by removing the need for standard curves, they are costly and may not be as accessible. We have also demonstrated that changes in RICE values can be validated at the bench without experimentally determining the actual RICE values. This can be done using a traditional qPCR approach without standard curves by designing qPCR experiments with the chimeric junction as the target and using the upstream/downstream parental regions as the endogenous control to calculate 5’ and 3’ template normalized ΔΔCt values for relative chimera expression. In this way, the qPCR result, like a RICE value, reflects the proportion of chimeric transcripts out of all transcripts containing the parent-chimera homologous region.

The value of differential RICE analysis with the FusionBlaster pipeline is demonstrated in this study through our investigation of prostate cancer chimeric RNAs. Here, we show that primary and metastatic prostate cancer sample cohorts processed at different times and by different groups cluster together and away from their respective noncancer tissue controls based on the RICE profile of approximately 1200 chimeric RNAs. We found a core set of 331 chimeric RNAs with significantly different RICE values common in primary prostate cancer, metastatic prostate cancer in liver, and metastatic prostate cancer in lung when compared with their respective GTEx tissue controls. Of the core set of 331 prostate cancer chimeric RNAs, 28 chimeras had significantly upregulated 5’ and 3’ RICE values and may serve as potential biomarkers or therapeutic targets for prostate cancer patients. Among the 28 chimeras with significantly upregulated 5’ and 3’ RICE values, we found the TE chimera, which results from the gene fusion of TMPRSS2 and ERG genes and is well established in the literature as the predominant molecular subtype of prostate cancer, validating the effectiveness of differential RICE analysis with FusionBlaster. Furthermore, we found several other chimeras that appear to have more apparent significantly upregulated RICE values between cancer groups compared to noncancer GTEx controls than the TE chimera.

Downstream gene set enrichment analysis of the parental genes of differentially expressed chimeric RNAs revealed a significant association with actin mediated cell contraction and membrane organization processes via the actinomyosin cellular component. Chimeric transcripts with parental genes EMP2, DSC2, DSP, MYL6B, PDE4D, and CTNNA3 were found to be differentially expressed in prostate cancer samples and disruption of their protein coding or regulation by chimeric transcript generating mechanisms may contribute to cytoskeleton remodeling processes in prostate cancer progression and metastasis. EMP2 for example, the parental gene of differentially expressed prostate cancer chimeras ATG10-EMP2, EMP2-FAM179A, and EMP2-SMAD2 found by the FusionBlaster pipeline, is known to contribute to prostate cancer metastasis (39–41) and controls cell migration, cell contraction, and focal adhesion density through its interactions with PTK2 and SRC (42–44).

RNA binding protein motif enrichment analysis was performed on sequences found upstream and downstream of chimeric RNA splice junctions resulting in RBPs that have established oncogenic roles. Furthermore, ELAV1/HUR, SRSF1, and PFS/SFPQ have been specifically reported to have implications in prostate cancer (36–38). HUR, for example, has been shown to translocate from the nucleus to the cytoplasm where it stabilizes mRNA transcripts, altering ERK5 activation to drive progression and metastasis in castration resistant prostate cancer (45, 46). Hur has been found to regulate mRNA stability and alternative splicing of glutaminase (GLS) transcripts leading to glutamine dependence in cancer cells (47). Recently, a large multiomics analysis on Chinese prostate cancer patients revealed SRSF1 expression and function as an essential oncogenic driver (48). PFS/SFPQ is responsible for alternate splicing and mRNA stability changes in androgen receptor (AR) transcripts driving proliferation in castration resistant prostate cancer cell lines and siRNA knockdown of PSF/SFPQ inhibited tumor growth of treatment resistant prostate cancer in mice (37).

In summary, the FusionBlaster pipeline provides a novel way to investigate chimeric RNAs by measuring the relative index of chimeric expression (RICE). The resulting RICE values accurately capture changes in chimeric RNA generation and can be validated in the laboratory. While FusionBlaster is not designed to detect novel chimeric RNAs, RICE values reveal differentially expressed chimeric RNAs between samples in a manner unbiased by read length and depth allowing the comparisons of different sequencing cohorts. The FusionBlaster pipeline is easy to install and run on a personal laptop. Using a single core and less than 100MB of RAM, 100 chimeric RNAs can be quantitated in approximately 30 minutes per sample. Using the FusionBlaster pipeline for differential RICE analysis in prostate cancer found TMPRSS2-ERG, a well-defined prostate cancer gene fusion, along with 27 other chimeric RNAs including GTSE1-TRMU, which has not been previously associated with clinical prostate cancer. Additionally, downstream analysis of chimeric RNAs with significantly different RICE values confirmed the involvement of cellular processes, components, and RBPS known to drive prostate cancer progression and metastasis while revealing novel insight on their implication with chimeric RNA expression.

## Supplemental Figure Legends

**Supplemental Table 1.**
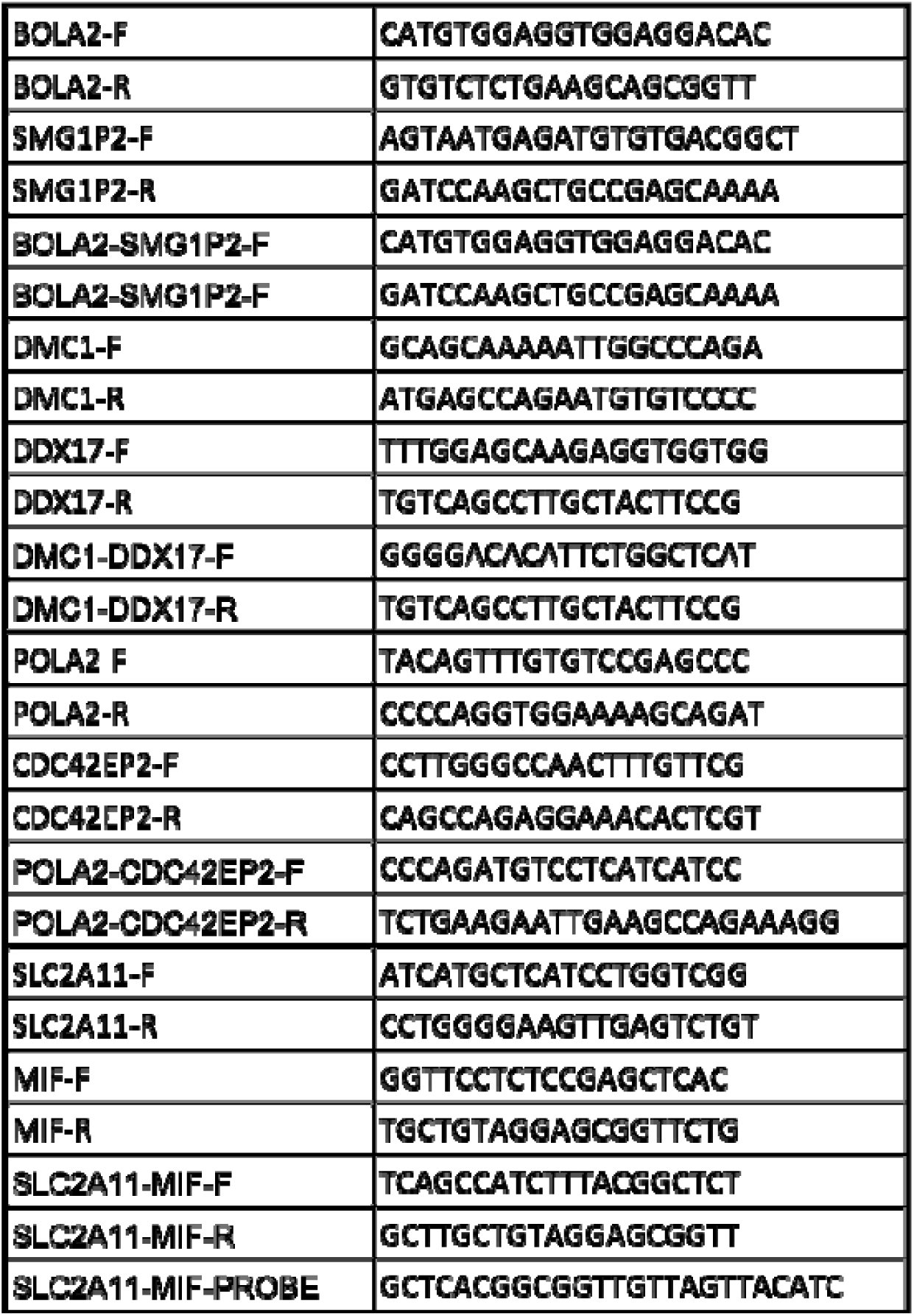
Primers used for RT-qPCR validation.

