## supplementary files for "Relative Index of Chimeric Expression (RICE) Analysis: A Quantitative Approach for Chimeric RNAs Using *FusionBlaster*"

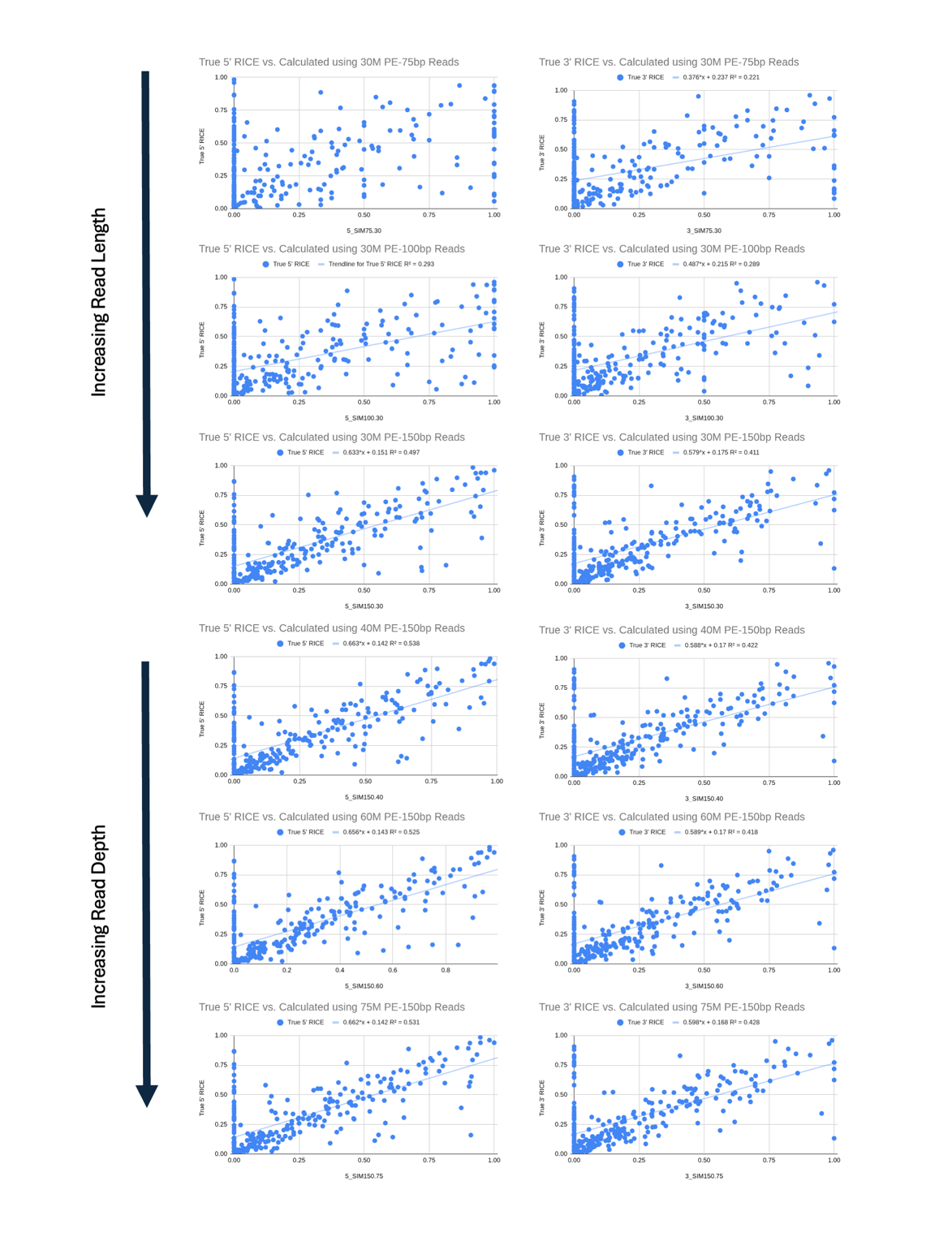

Supplemental Figure 1- STAR-based correlation with True RICE values

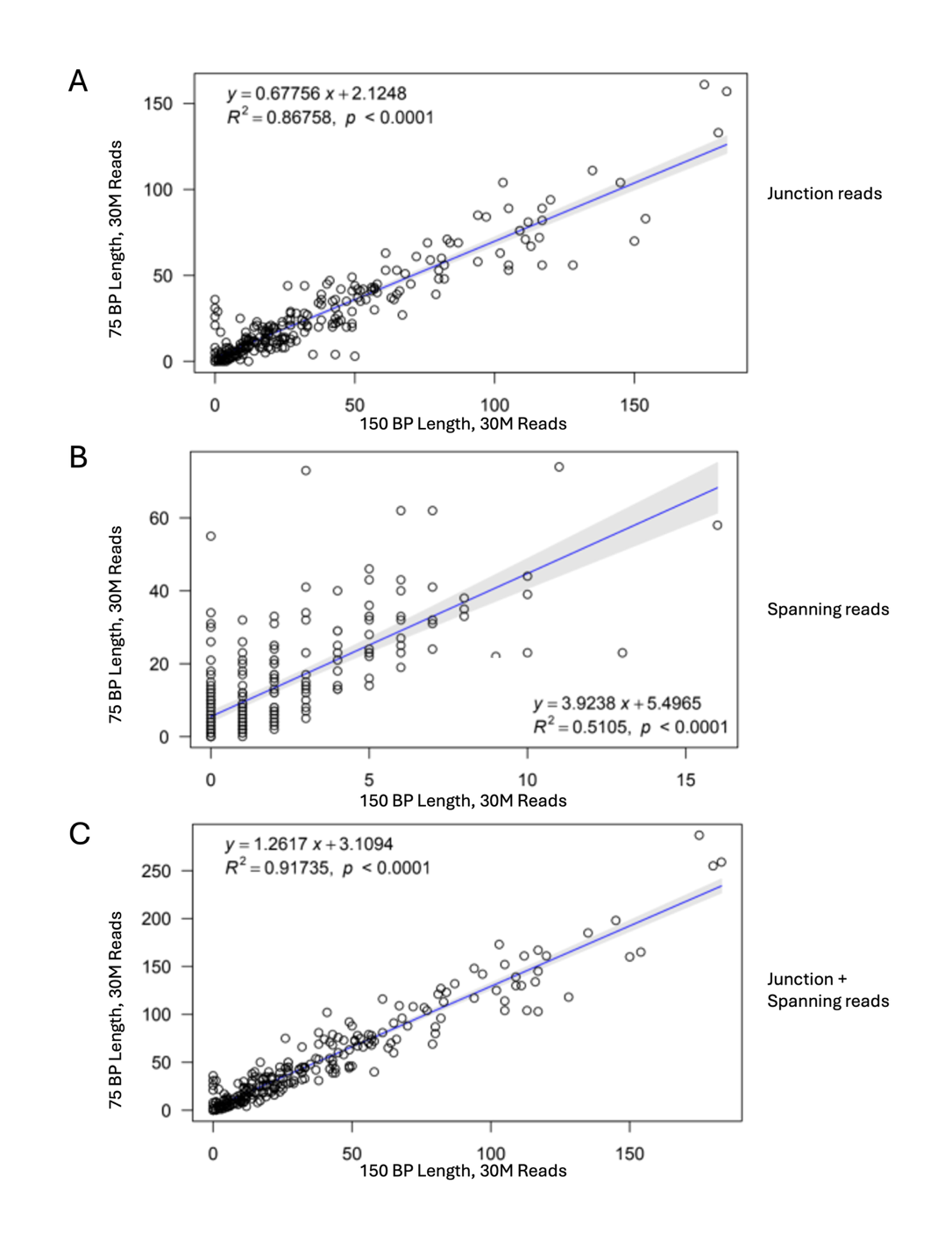

Supplemental Figure 2- STAR-based correlation with True RICE values

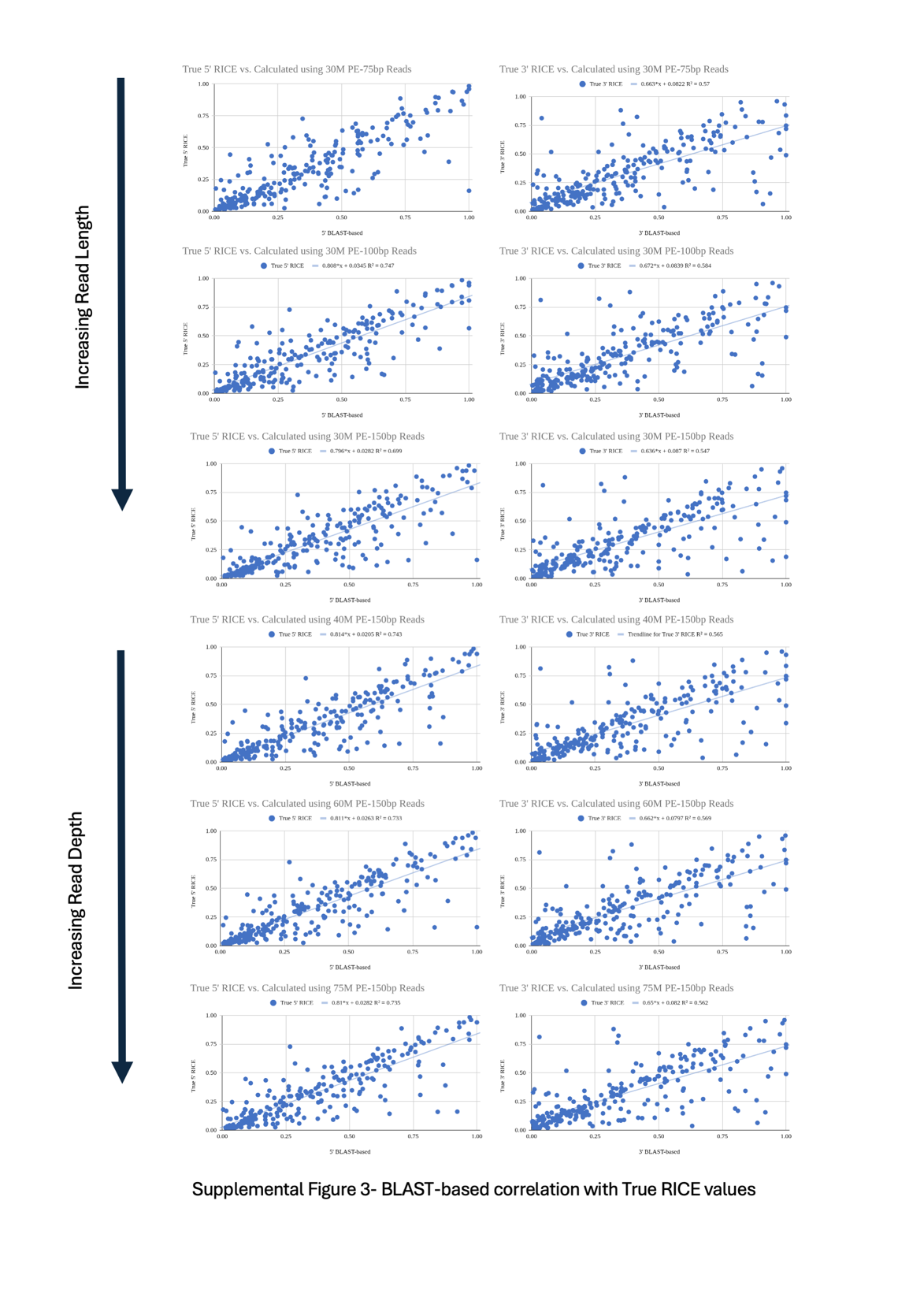

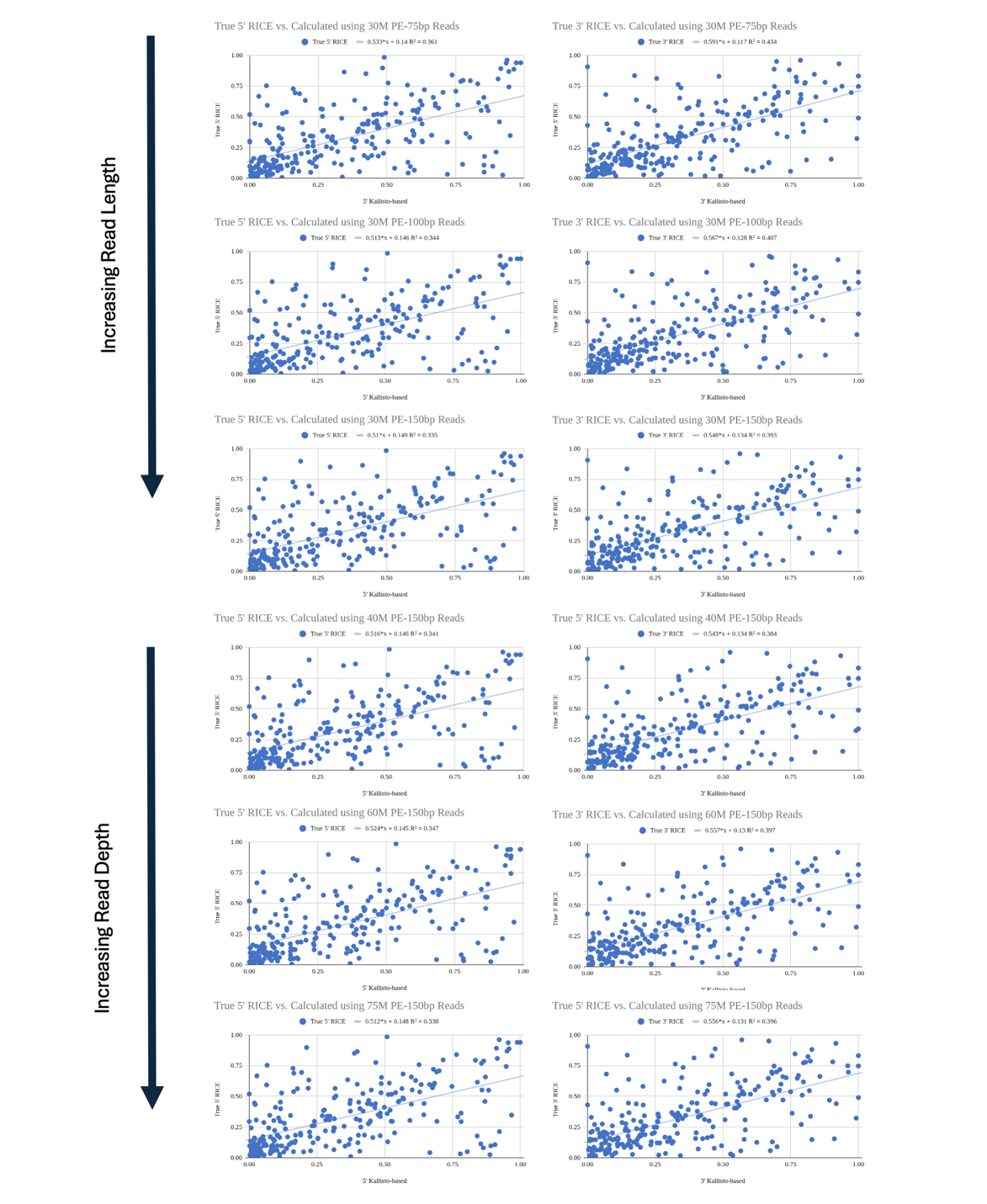

Supplemental Figure 4- Kallisto-based correlation with True RICE values
